# Clonal thermal preferences affect the strength of the temperature-size rule

**DOI:** 10.1101/2020.04.08.031286

**Authors:** Anna Stuczyńska, Mateusz Sobczyk, Edyta Fiałkowska, Wioleta Kocerba-Soroka, Agnieszka Pajdak-Stós, Joanna Starzycka, Aleksandra Walczyńska

## Abstract

Genetically similar organisms act as a powerful study system for the subtle differences in various aspects of life histories. The issue of trade-offs among traits is of special interest. We used six parthenogenetic rotifer clones previously exposed to different thermal laboratory conditions. Interclonal differences in female body size were examined in common garden conditions. We estimated the population growth rate and strength of the size-to-temperature response across four thermal regimes. We tested hypotheses on the existence of the relationships between (i) thermal acclimation and species body size, (ii) thermal specialization and fitness and (iii) thermal specialization and strength of the temperature-size rule. Positive verification of (i) would make it justifiable to refer the other investigated traits to thermal preference and, further, to thermal specialization. Addressing the issues (ii) and (iii) is our pioneering contribution to the question on the strength of size-to-temperature response as differing across life strategies. We hypothesized that this plastic response may be affected by the level of thermal specialization and that this pattern may be traded off with the temperature-dependent potential for population growth rate. Additionally, we investigated the differences in reproductive strategy (number of eggs laid by a female and female lifetime duration) in one temperature assumed optimal, which acts as an important supplement to the general clonal life strategy. We confirmed that the thermal acclimation of a clone is related to body size, with clones acclimated to higher temperatures being smaller. We also found that warm-acclimated clones have a narrower thermal range (= are more specialized), and that the temperature-size rule is stronger in rotifers acclimated to intermediate thermal conditions than in specialists. Our results contribute into the issue of trade-offs between generalist and specialist strategies, in the context of plastic body size respone to different temperatures.

## Introduction

Optimal allocation theory states that living organisms optimize their strategies of the allocation of limited resources among the functions associated with maintenance, repair and reproduction. Any given set of abiotic and biotic conditions provides a unique, optimal strategy, and trade-offs among life history traits are at the base of such strategies (Cody 1966; Kozłowski 1992; Kozłowski 2006; Roff 1992; Stearns 1992). The pattern of limited resources causing the traits to trade is complex and troublesome with regard to straightforward inferences and conclusions (Acerenza 2016; Lailvaux and Husak 2017; Roff and Fairbairn 2007). Going a step further, DeWitt (2016) stressed the importance of including not only trait means but also the shape (i.e., skew) of phenotypic plasticity in studies on life-history optimization in ecological contexts. According to the author, the strength of phenotype distribution across environments should be optimized by means of natural selection. Phenotypic plasticity is a non-random variance in a given trait that does not involve differences in genotype, an idea that has been well established since Levins (1968). However, this idea is still not well integrated with life-history theory (DeWitt 2016). In this study, we investigated how the strength of the phenotypic size-to-temperature response varies in six parthenogenetic rotifer clones previously exposed to different temperatures in the laboratory.

The phenotypic achievement of smaller body size with increasing temperature in ectotherms is a widely observable but not fully understood phenomenon called the temperature-size rule (TSR, Atkinson 1994). Revealing the adaptive significance of this pattern is one of the largest challenges facing evolutionary biology (Berrigan and Charnov 1994), especially given that it concerns one of the most important organismal traits, body size (Hildrew et al. 2007; Kozłowski 2006; Stearns 1992), and one of the most evolutionarily influential environmental variables, temperature (Johnston and Bennett 2008). Direct confirmation of the adaptive significance of the TSR was provided by Walczyńska et al. (2015a) for *Lecane inermis* rotifer. In the first step of our study focusing on the same species, our goal was to confirm whether thermal acclimation is linked with thermal preference through body size at the interclonal level, namely, whether the clones prefer the temperature they were acclimated to. Given that the TSR is the adaptive body size adjustment to given thermal conditions, clones acclimated to lower/higher temperatures should prefer the respective thermal range, and further, those acclimated to lower/higher temperature should display larger/smaller body sizes, respectively.

The next step of the study was to link thermal preference with thermal specialization through fitness. We predicted that clones preferring low or high temperature would specialize to the respective thermal range and would perform better (= have higher fitness) at those respective temperatures than other clones. This prediction is based on the optimal allocation theory, which assumes the existence of resource limitations that force trade-offs and consequently diversify the life strategies of organisms. Some historical predictions were made regarding the level of ecological specialization and performance. The most classical one was proposed by Levins (1968). He assumed a distinction between generalists, characterized by relatively high and flat performance across environments, and specialists, which perform better in one type of environment than in others. This idea was challenged multiple times, and the thermal biology field displays both empirical evidence in favor of (Foray et al. 2011; Griffith and Sultan 2006; Verberk et al. 2010; Walczyńska and Serra 2014) and against predictions (Berger et al. 2014; Knies et al. 2009; Sheth and Angert 2014). However, the most intuitive argument in favor of the existence of such a trade-off came from work on the temperature dependence of proteins (Fields 2001), which showed that enzymes can be either molecularly stable (a “specialist”) or structurally flexible (a “generalist”) depending on the physical role they play, but not both. Therefore, enzymes adapted to work at one temperature will be unable to function at suboptimal temperatures unless their structure is modified (Fields 2001).

In the final step, we switched from the single phenotype to the variable phenotypes perspective, namely, phenotypic plasticity. Divergent natural selection resulting from environmental variation creates functional trade-offs between environments; organisms adapted to one environment present reduced performance in alternative environments (DeWitt and Langerhans 2004). Moreover, theoretical modeling predicts that the strategy of intermediate phenotypes, the generalists, is to reduce variance in performance across environments (DeWitt and Langerhans 2004). We use this theoretical background for the first time in the context of the strength of plastic size response to temperature. We test this novel suggestion on six clones of *Lecane inermis* rotifer species.

Under these theoretical assumptions, we aimed to match the general interclonal potential for phenotypic plasticity in size response to temperature with performance measured in the most universal evolutionary currency, fitness. To date, there is no information in the literature on the possible trading off between the size-to-temperature response and general thermal preferences, except for some suggestions presented by Walczyńska and Serra (2014). The authors suggested that the differences in the strength of the TSR response found in three *Brachionus* species belonging to the *Brachionus plicatilis* cryptic species complex might be attributed to the level of species thermal specialization, with generalists showing more flexible plastic body size adjustment across temperature than specialists (Walczyńska and Serra 2014).

Clones, which are highly genetically similar organisms, provide a promising system for understanding the subtle details behind the observed eco-evolutionary processes. The interclonal perspective in studies on the temperature effect was previously successfully applied to reproductive strategy in the *Lecane inermis* rotifer studied here (Fiałkowska et al. 2011) and to size at maturity and asymptotic size (Hoefnagel et al. 2018) and vital rates (Bruijning et al 2018) in *Daphnia magna*. In this study, we explored the unique opportunity of possessing in the laboratory six clones of *Lecane inermis* Bryce (Rotifera) that were previously exposed to different thermal regimes. We assumed that they exemplified different thermal preferences gained through adaptive mechanisms. Our reasoning involved treating the clone-specific body size information as a link between thermal acclimation (warm-, intermediate temperature- or cold-acclimated clones) and thermal preference, based on the theoretical predictions as described above. If our predictions are correct, this linkage would justify that we refer our results to the clonal thermal preferences. We estimated the thermal sensitivity of both the population growth rate *r* and the size-to-temperature response across the same four thermal regimes. Additionally, we searched for possible differences in the clone-specific reproductive strategy of females (number of eggs laid and duration of life) at the temperature assumed optimal. Such differences would facilitate the interpretation of the results on the general performance and the abilities of plastic response to environmental changes.

Based on the theoretical predictions on the performance of generalists *vs*. specialists in different environments (DeWitt and Langerhans, 2004; Levins, 1968) we hypothesized the following patterns in phenotypic size-to temperature response (Atkinson, 1994):

H1: body size of clones differs according to their thermal acclimation; cold-acclimated clones are the largest, followed by intermediate clones, and the smallest are warm-acclimated clones;

H2: cold- and warm-acclimated clones are more specialized (= have higher fitness) to lower or higher temperatures, respectively, than clones acclimated to intermediate temperatures (= generalists);

H3: phenotypic size response to temperature (TSR) is stronger in clones acclimated to intermediate temperatures than in more specialized cold-acclimated and warm-acclimated clones.

## Material and Methods

### Clones isolation and maintenance

*Lecane inermis* is a bacterivorous, monogonont rotifer species occurring in psammolittoral fresh and brackish water bodies that is relatively common worldwide (Bielańska-Grajner et al. 2015). It is a frequent inhabitant of wastewater treatment plants (Klimowicz 1989), from which all the investigated clones were isolated. Its lifecycle consists of the sexual and asexual phases (Miller 1931), although in most cases, the sexual phase disappears in the populations from the wastewater treatment plants (Pajdak-Stós et al. 2014). The generation time estimated for one clone is approximately 2 days between 15 °C and 25 °C (Walczyńska et al. 2015b), while a doubling time for another four *L. inermis* clones was estimated to be 1.48-1.73 days at 20 °C (Fiałkowska et al. 2011). The size of eggs constitutes the exceptionally large proportion of the female size, and consequently, the size of newly hatched females is 71% of adult females on average (Miller 1931). *L. inermis* has been intensively investigated regarding the phenomenon of the temperature-size rule. It follows this rule both in the laboratory and in the field (Kiełbasa et al. 2014) and displays adaptive body size adjusted to temperature-dependent oxygen levels (Walczyńska et al. 2015a) in a mechanism controlled at two points within the lifecycle (Walczyńska et al. 2015b).

Details about the isolation and laboratory maintenance of the clones used in the study are provided in Table 1. For each clone, the isolation was started from one individual fed with 25 μl of 3‰ molasses solution or 10 μl of suspension of NOVO, a nutrition powder used for rotifer mass culture (patented by Pajdak-Stós et al. 2017), 0.20 g NOVO in 50 ml of Żywiec spring water. The proliferated clones were cultured in 24-well tissue culture plates in Żywiec brand spring water (Poland), fed with NOVO and kept in darkness. Clones selected for high performance at a lower temperature, specifically 8 °C or 15 °C, were transferred to the new wells in the tissue culture plates every two weeks. Other clones were passaged every week. The clones included in Table 1 are those showing the highest proliferation under such selection from among all the clones tested. In 09/2016, all the clones, except for Warm2, were transferred to 25 °C (darkness) in Żywiec medium-fed NOVO. We assumed that the best-performing clones underwent some adaptive mechanisms to the given thermal conditions, and this condition-specific selective force might have differentiated the clonal general life histories. Clone Warm2 was transferred from 20 °C to 25 °C in 06/2017. All clones were exclusively parthenogenetic, which is typical of *L. inermis* from this type of habitat (Pajdak-Stós et al. 2014).

**Table 1.** Description of the origin and the thermal conditions of the maintenance (= acclimation) preceding this study for the experimental clones of *Lecane inermis* rotifer. WWTP – wastewater treatment plant

| code | description | thermal preferences |
| --- | --- | --- |
| **Int1** | isolated in WWTP1 in Poland in 2014, maintained at 15 °C | intermediate |
| **Int2** | isolated in WWTP1 in Poland in 2014, maintained at 15 °C | intermediate |
| **Int3** | isolated in WWTP1 in Poland in 2014, maintained at 15 °C | intermediate |
| **Warm1** | isolated in WWTP2 in Poland in 2010, maintained at 20 °C | warm<sup>a</sup> |
| **Warm2** | isolated in WWTP near Barcelona, Spain in 2011, maintained at 20 °C | warm |
| **Cold** | isolated in WWTP3 in Poland in 2015, maintained at 8-10 °C for 10 months, later transferred to 15 °C | cold |

<sup>a</sup>optimal temperature estimated at 31 °C by Walczyńska et al. (2016)

### Body and egg size

For each clone, the subsamples were taken from the stock populations maintained in common garden conditions at 25 °C and were fixed with Lugol solution. Approximately 50 females and 30 eggs per temperature were photographed using the Nikon Eclipse 80i microscope, Nikon DS-U1 camera and NIS Elements software. The length and width were measured in ImageJ (NIH, USA), and their product was treated as an area measure.

### Fecundity

In 06/2017, 60-mm Ø Petri dishes with a small number of females representing three clones, Int2, Cold, and Warm2 (one dish per clone), were transferred to 30 °C and fed with 10 μl of the commercial bioproduct Biotrakt^®^ (Zielone oczyszczalnie, Poland). The choice of temperature was dictated by the results indicating that the optimal temperature for *Lecane inermis* clone Lk6 was 31 °C (Walczyńska et al. 2016). After two days, which was necessary to obtain the next-generation females from freshly laid eggs, 24 individual young females per clone were transferred to separate wells in 24-well tissue culture plates (TPP, Switzerland) with 1 mL of Żywiec medium and 10 μl of 10× diluted Biotrakt^®^ per well. The number of eggs laid per female was checked daily under a stereomicroscope. In one case, the gap between two checks was two days. If eggs were observed, a female was transferred to a new well (with the same conditions as previously described) to avoid the counting of eggs laid by the next-generation females. The death of each female was noted. This procedure was continued until the last female died. After completion of this stage, the same procedure was conducted for another three clones: Int1, Int3 and Warm1. The eggs laying was examined daily except for one case, where the gap between the inspections was two days.

### Growth rate estimation

In 07/2017, populations of all six clones were established in four temperature treatments: 15, 20, 25 and 30 °C. The initial numbers were 10 females × three replicates per clone. The cultures were kept in six-well tissue culture plates (TPP, Switzerland), with Żywiec as the medium supplemented with 20 μL of Biotrakt^®^. The number of females was counted twice, with a 1-2-day gap between the counts, to correct for growing processes being physically dependent on temperature (a longer gap was applied at the two lower temperature regimes). These numbers were then used to calculate the population growth rate *r* according to the formula *r* = (ln(*x_2_*) - ln(*x_1_*))/*t*, where *x_2_* – number of individuals in count II, *x_1_* – number of individuals in count I, and *t* – time in days.

### Size-to-temperature response examination

In 08/2017, approximately 60 individuals per clone per temperature, derived from the cultures maintained for growth rate estimation, were fixed with Lugol solution for size measurements. The rotifers were photographed using a Nikon Eclipse 80i microscope, Nikon DS-U1 camera and NIS Elements software. The lengths and widths of individuals were measured using ImageJ (NIH, USA), and their product was used as a proxy for area.

### Statistical analyses

All the analyses were performed in Statistica 13 (StatSoft 2014). Parametric tests were conducted when the assumptions were met. Under assumptions violation, the non-parametric tests were conducted. In the case of size, simple regression was used to test for the relationship between clone-specific female size and egg size. We tested for the differences in the number of eggs laid per day of female life using the Kruskal-Wallis test. The differences in the growth rate between clones were tested using one-way ANOVA for each temperature separately (the lack of homogeneity of variance prevented joint analysis). The differences in body size between clones (the plastic size responses) were tested using one-way ANOVA for each temperature separately because of the violation of the assumption of the homogeneity of variance. The same reason prevented parametric analysis at 30 °C. In this case, the differences were tested with the Kruskal-Wallis test. To compare the temperature effect on growth rates among the groups of thermal acclimation, one value describing a common slope of response across temperature was estimated for each clone using the simple regression model (statistics presented in Table 2), and then these values were compared among clones in a one-way ANOVA for the groups of warm-, cold- and medium-temperature acclimation. The same procedure was applied to compare the strength of TSR among the groups of thermal specialization, and the statistics are presented in Table 2.

**Table 2.** Statistics for the estimation of the slopes for population growth rate and size across temperature.

| trait | Clone | slope | $R^2_{(adj)}$ | t (df) | p |
| --- | --- | --- | --- | --- | --- |
| population growth rate | Cold | 0.083 | 0.65 | 4.63 (10) | 0.00093 |
|  | Int1 | 0.068 | 0.82 | 7.07 (10) | 0.00003 |
|  | Int2 | 0.041 | 0.69 | 5.03 (10) | 0.00051 |
|  | Int3 | 0.028 | 0.29 | 2.27 (9) | 0.04964 |
|  | Warm1 | 0.084 | 0.79 | 6.42 (10) | 0.00008 |
|  | Warm2 | 0.053 | 0.71 | 5.22 (10) | 0.00039 |
| TSR | Cold | -0.021 | 0.33 | -8.33 (140) | < 0.000001 |
|  | Int1 | -0.022 | 0.43 | -9.09 (108) | < 0.000001 |
|  | Int2 | -0.025 | 0.52 | -11.39 (117) | < 0.000001 |
|  | Int3 | -0.023 | 0.29 | -6.52 (101) | < 0.000001 |
|  | Warm1 | -0.015 | 0.27 | -7.23 (136) | < 0.000001 |
|  | Warm2 | -0.014 | 0.22 | -5.75 (111) | < 0.000001 |

## Results

### Body and egg size

The female size in μm^2^ and respective egg size for each clone are provided in Table 3. There was no interclonal relationship between female size and egg size (slope = 0.15, R^2^_(adj)_ = 0.025, p = 0.35; Fig. 1A). The average egg size as a proportion of mother size reached 78% (from 73% for Int3 to 85% for Warm1).

**Table 3.** The mean body and egg size for studied clones for individuals sampled from the common garden stock at 25 °C. Ordered from the smallest to the largest clone.

| clone | Female size ($\mu\text{m}^2$) | Egg size ($\mu\text{m}^2$) |
| --- | --- | --- |
| | mean ± SE; N | mean ± SE; N |
| **Warm1** | 2517 ± 32; 55 | 2147 ± 25; 30 |
| **Warm2** | 2665 ± 46; 55 | 2098 ± 25; 29 |
| **Int2** | 2799 ± 40; 55 | 2201 ± 28; 30 |
| **Int1** | 2812 ± 42; 54 | 2078 ± 34; 29 |
| **Cold** | 2848 ± 56; 54 | 2182 ± 24; 30 |
| **Int3** | 3019 ± 54; 53 | 2210 ± 33; 24 |

**Fig. 1.**
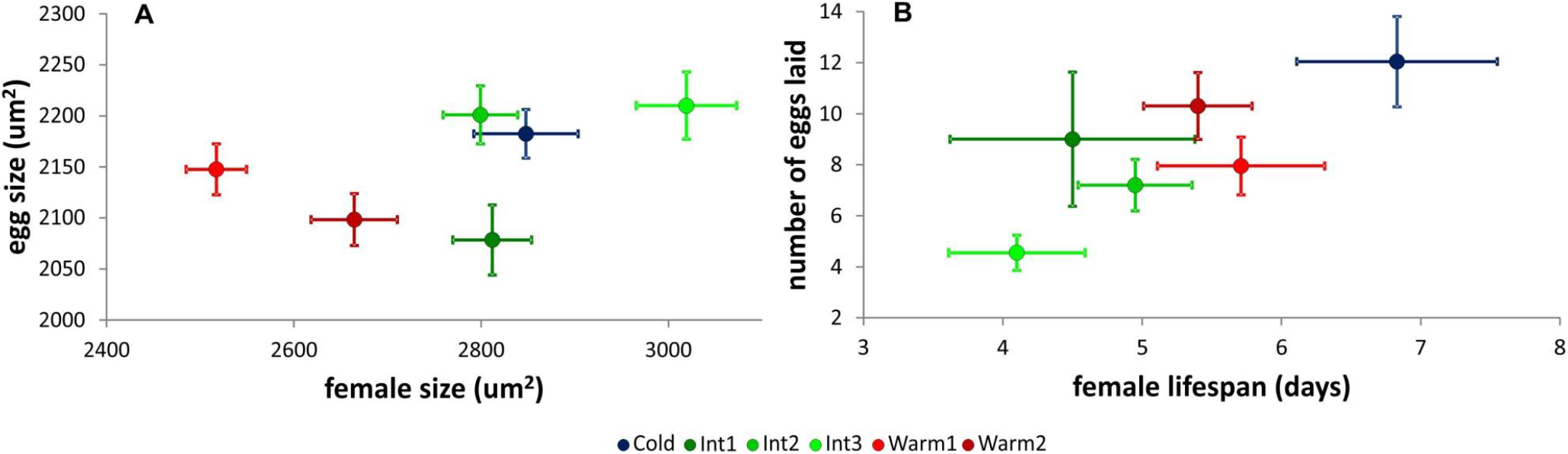
Relationships among life history traits for six clones of *Lecane inermis* rotifer: female body size and egg size measured from the stock populations at 25 °C (A), female lifespan and total number of eggs laid per female estimated at the temperature of 30 °C assumed optimal (B). Thermal adaptation distinguished by color. Means ± SE

### Fecundity and lifespan

At the common temperature of 30 °C assumed optimal, clones differed in the mean number of eggs laid by the females (Kruskal-Wallis test: H_(5, 125)_=15.3, p = 0.009) and in the female lifespan duration (Kruskal-Wallis test: H_(5, 125)_=13.9, p = 0.016). The females laid significantly more eggs with the lifetime duration (slope = 2.2, R^2^_(adj)_ = 0.76, p < 0.0001), with the Cold clone being the most fecund, followed by both warm clones and, finally, the least fecund intermediate clones (Fig. 1B).

### Growth rate and temperature-size rule

There were significant differences among clones in growth rate at each temperature. According to the Tukey post hoc tests,

- 15 °C, Int2 clone had significantly faster growth than clone Warm2, while the other four clones showed intermediate patterns;
- 20 °C, clones Int2 and Warm2 grew significantly faster than clone Int3;
- 25 °C, Cold clone performed significantly better than clones Int2, Int3 and Warm1;
- 30 °C, clone Warm1 grew faster than clones Warm2 and Int3, and clones Cold and Int1 performed better than clone Int3 (Fig. 2A).

**Fig. 2.**
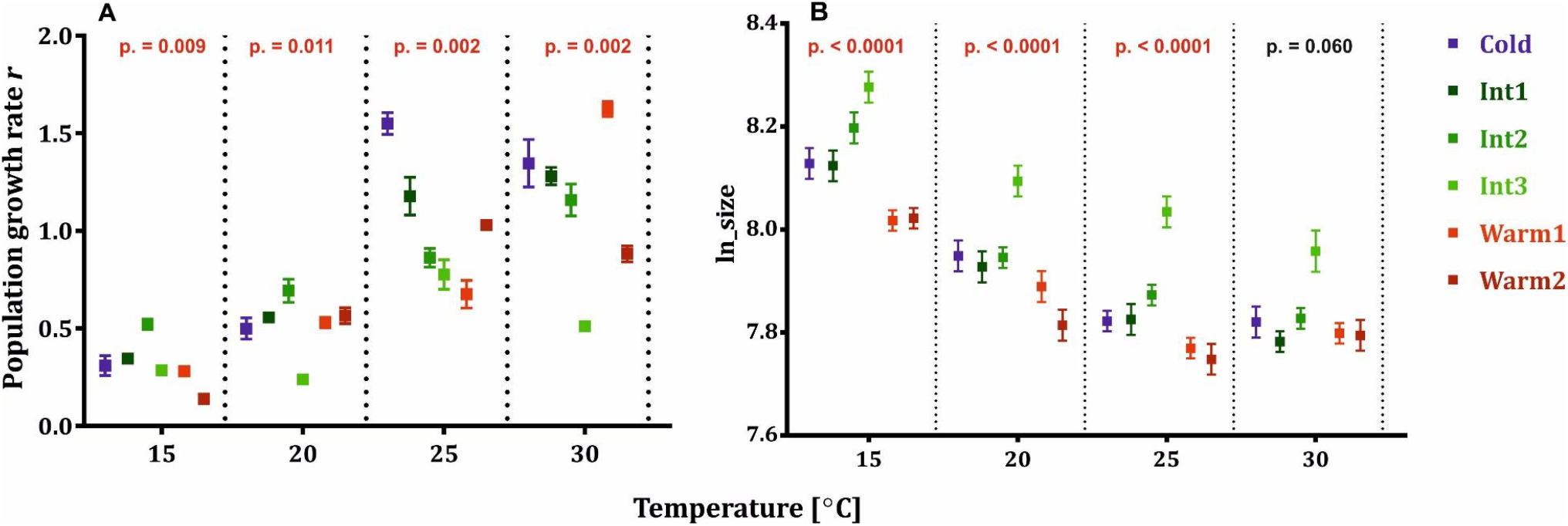
Interclonal response to temperature in six clones of *Lecane inermis* rotifer: population growth rate *r* (A) and temperature-size rule response (B). Means ± SE.

All rotifers were smaller with higher temperature (slope = – 0.11, R^2^_(adj)_ = 0.34, p < 0.001), and significant differences among clones existed in all but the 30 °C treatment (Fig. 2B). Generally, clone Int3 was the largest across the regimes, while the smallest were both warm clones. The Cold clone presented an intermediate pattern together with the remaining two Int clones.

The analysis of the across-temperature ability to proliferate and to plastically respond (Table 2) showed the opposite effect of temperature. The interclonal comparison of these two temperature-dependent effects (with the strength of the size-to-temperature response presented as a positive effect for the sake of clarity), with the clones pooled within the thermal acclimation groups (N = 3), showed a general lack of relationship (slope = −0.087, R^2^ = 0.198, p = 0.38; Fig. 3). The population growth rate was not different across specialization (F_(2,3)_ = 1.39, p = 0.37; Fig. 3, x-axis), while a significant difference occurred in the case of the strength of TSR across groups (F_(2,3)_ = 27.53, p = 0.012; Fig. 3, y-axis). Clones acclimated to intermediate temperature displayed a significantly stronger (according to Tukey’s HSD test) size-to-temperature response than warm-acclimated clones, while Cold clone represented an intermediate pattern.

**Fig. 3.**
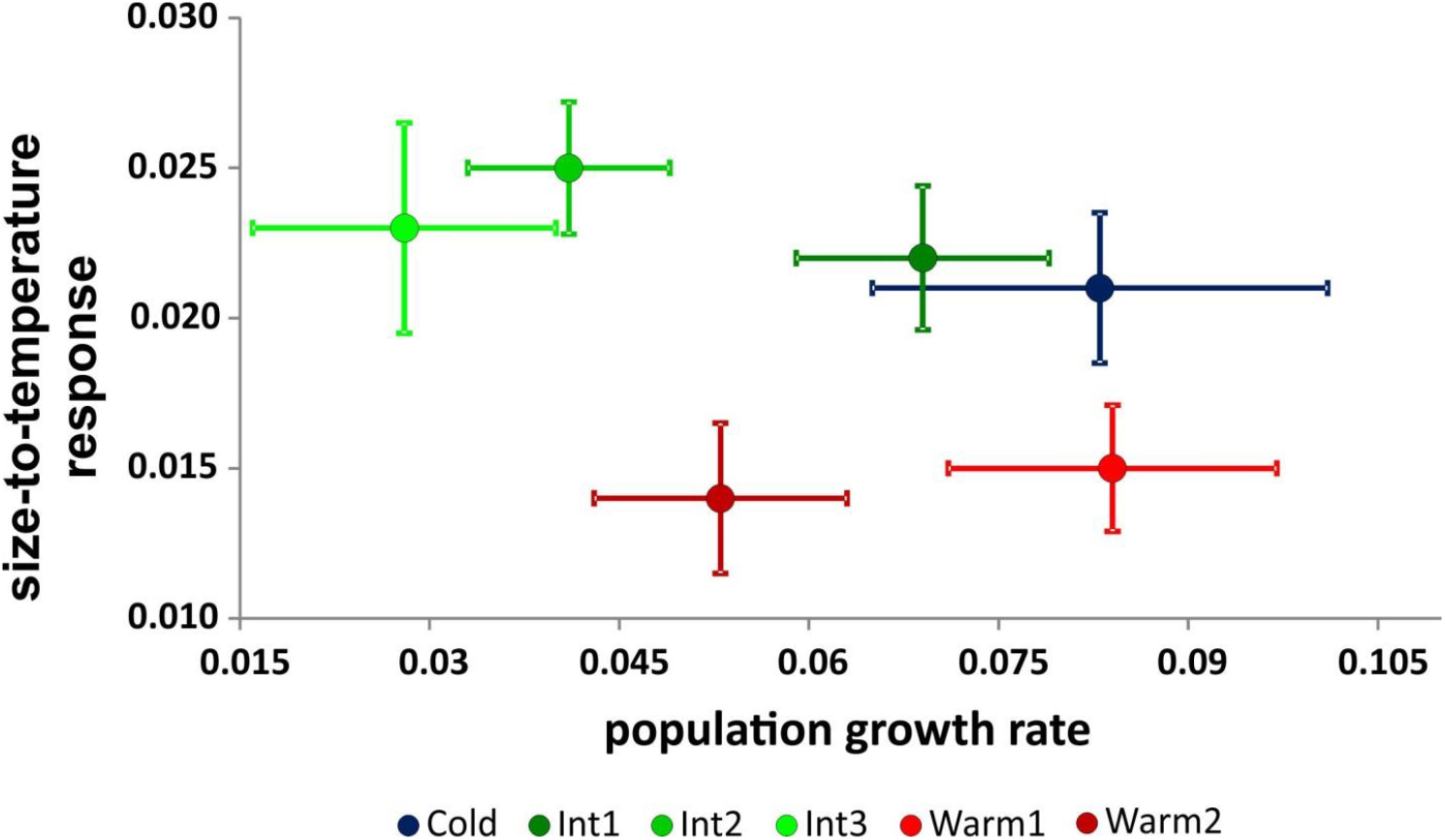
Relationship between the slopes of the following: increase in population growth rate *r* (x) and of body size decrease (the absolute values of the negative slopes; y) across temperature. Each point represents a clone with its thermal adaptation distinguished by color. slope ± SE

## Discussion

In this study, we present the interclonal link between thermal preference, ability of plastic response and fitness to elucidate the issue of ecological trade-offs. First, in hypothesis H1, we tested whether thermal acclimation could be referred to as thermal preference in the six investigated clones. According to our predictions, the cold-acclimated clone was larger than the two warm-acclimated clones (Fig. 1A). The three clones acclimated to intermediate temperature represented a wide range of sizes, although they were consistently larger across temperatures than warm-acclimated clones (Fig. 1A), also in accordance with our predictions. We expected the Cold clone to be the largest, which was not the case; however, based on the size measurements from the stock populations, only one Int clone appeared larger than the Cold clone (Fig. 1A). Therefore, we generally validated our assumption that it is justified to refer to thermal preference instead of thermal acclimation in our further interpretation of the results. Additionally, this result confirms the adaptive nature of body size linked to temperature at the interclonal level along with the previous interspecific patterns found for three species representing the *Brachionus plicatilis* cryptic species complex (Walczyńska and Serra 2014) and five species of free-living nematodes (Majdi et al. 2019).

Regarding hypothesis H2 on the relationship between thermal preference and fitness across temperature, warm clones were the poorest performers at cold temperatures and were considerably better at higher temperatures, especially Warm1 (Warm2 appeared to have a thermal optimum closer to 25 °C than to 30 °C; Fig. 2A). The cold-acclimated clone performed notably well at high temperatures but was an intermediate performer at the lowest temperature. Int-clones showed different patterns, with Int1 and Int2 performing relatively well at all temperatures (a clear generalist strategy) and Int3 performing poorly in all thermal regimes (Fig. 2A). Our results suggest that specialization is restricted to warm-acclimated organisms rather than to mid-temperature-acclimated organisms, which present the strategy of generalists, thus partly confirming our predictions in H2. The case of the Cold clone is more complicated. At the lowest temperature, it had higher fitness than warm-acclimated clones but similar or lower fitness than Int clones. However, it is important to note that the lowest experimental temperature was not the one to which a Cold clone was acclimated, and using a temperature below 15 °C might result in the Cold clone displaying higher fitness than other clones, the prediction supported by the pilot study presented in the supplementary materials (Fig. S1). Such a thermal regime was not planned in the study because it was previously reported that the minimum temperature for the proliferation of a clone referred to here as Warm1 was 11 °C (Walczyńska et al. 2016).

Therefore, we set the lowest thermal regime to 15 °C to compare the positive proliferation of all six clones. The result we obtained for a Cold clone confirms the classical models in which organisms acclimated to the use of a wider range of resources perform better (originally, they outcompete) those of a narrower range (Tilman 1982). Our Cold clone was acclimated to the usage of resources at the lowest thermal regimes (down to 8 °C), unlike all the others. This result is more in keeping with the findings described by Zamorano et al. (2017), with cold-preferring organisms performing notably well in the whole range studied, than with the findings reported by Majdi et al. (2019), with the largest species performing well only in the coldest regime. Perhaps there are other factors that cause the discrepancies among these studies; one of the most possible factors is body size. Our results on clone-specific body size show that it is better not to be the largest but rather to have an intermediate size, unless the temperature is high. In this way, we solved the conflict in the haiku in Kingsolver and Huey (2008): *Bigger is better. And hotter makes you smaller. Hotter is better*. The solution is that hotter is better, but only for smaller, a result previously achieved for the *L. inermis* rotifer (Walczyńska et al. 2015a). This result was obtained in the context of oxygen as a driving factor in the TSR, as theoretically assumed (Atkinson et al. 2006; Verberk et al. 2011), and provides one of the limited confirmations of the adaptive role of small size under a combination of high temperature and low oxygen availability. In this study, six clones of *Lecane inermis* rotifer displayed a consistent decrease in size with increasing temperature regardless of their thermal acclimation and the resulting thermal preferences. The relationship between the strength of this response and thermal specialization (hypothesis H3) showed that clones acclimated to medium temperature displayed a stronger decrease in size with increasing temperature than cold- and warm-specialized clones (Table 2, Fig. 2B). This pattern was not a compensation of size plasticity for the thermal sensitivity of the population growth rate because their relationship indicating a possible trade-off was not significant (Fig. 3). However, this result may be attributable to a small sample size, and the possibility of such compensation cannot be fully excluded. Both Cold and Warm clones displayed a weaker size-to-temperature response than Int clones, showing that the strength of this plastic reaction is not just a linear effect of acclimation temperature.

The thermal regimes were chosen to reflect the optimal thermal range previously invoked to designate the frames for TSR performance (Atkinson et al. 2003; Walczyńska et al. 2016). Thus, the responses we examined were not the result of exposure to stressful conditions on any end of thermal range, and they therefore did not deviate from the TSR in a classical sense. In addition, we link this pattern with thermal preferences and the level of thermal specialization. We show for the first time that organisms acclimated to intermediate temperature performed relatively well across temperatures, thus confirming the theoretical prediction that intermediate phenotypes as generalists (DeWitt and Langerhans 2004) display stronger TSR responses than specialists. This result presents the first empirical confirmation of the previous speculations made for three *Brachionus plicatilis* sister species (Walczyńska and Serra 2014).

The set of hypotheses stems from the theoretical predictions on generalist-specialist trade-offs in the context of adaptation to thermal conditions. Accordingly, the most beneficial temperature is expected to be the one to which an organism was acclimated (beneficial acclimation hypothesis, BAH; Leroi et al. 1994). Alternatively, the adaptation may refer only to low temperature acclimation (colder is better, CIB), hot acclimation (hotter is better, HIB), optimal (intermediate) temperature (optimal developmental temperature hypothesis), and other variants (reviewed in Deere and Chown 2006). However, none of the variants was found to be universal. Our results generally confirm the beneficial acclimation hypothesis (BAH) – clones exposed for a long time to specific thermal conditions display a preference (= perform better) for these conditions – but this effect is not consistent within a thermal continuum. The second-to-largest Cold clone was the best performer across the regimes, confirming the results found previously for parasitoids (Zamorano et al. 2017) and thereby confirming the colder-is-better hypothesis (CIB). On the other hand, Warm clones, the smallest, confirm the previous positive results for hotter-is-better (HIB) (Kingsolver 2009; Knies et al. 2009). The difference in these two clones is that Warm1 was previously reported to have a very high optimal temperature despite its origin from a temperate region, while Warm2 was assumed to be Warm-acclimated because of its origin from the Mediterranean region (Table 1). Interestingly, in light of our results, the Warm1 clone was more specialized at high temperatures than Warm2. Two mid-sized medium temperature clones performed relatively well and evenly in the thermal range studied, reflecting a clear generalist pattern, and a similar result was obtained for mid-sized nematode species (Majdi et al. 2019). Our largest clone, also acclimated to intermediate temperature, exhibited the poorest performance of all (though best at the lowest temperature as relative to other clones). Regarding nematodes, the largest species showed notably poor growth in all other regimes compared with the other four species, although it had the highest fitness in the coldest temperature (Majdi et al. 2019). These results are in contrast to previous reports (Geister and Fischer 2007; Kingsolver and Huey 2008) and show that larger sizes are not always better. Our novel contribution to the discussion of the superiority of the BAH, CIB or HIB hypotheses is that the answer is context-dependent, and that context is the level of thermal specialization of the investigated organisms.

The additional information on the reproductive traits shed light on the interclonal differences in general life strategies. In contrast to the optimal theory models (Fox and Czesak 2000; Honěk 1993; Parker and Begon 1986), we found no interclonal trade-off between female size and egg size (Fig. 1A) or between female lifespan and fecundity (Fig. 1B), though such trade-offs cannot be excluded at the level of individuals. The most apparent trade-off we found was allocation in reproduction *vs*. ability to proliferate across temperature between specialists and generalists. The cold-acclimated clone was the most fecund, and the two smallest, warm-acclimated clones displayed an intermediate pattern, while intermediate temperature-acclimated clones displaying the generalist strategy (evenly good performance at all temperatures) were the shortest-living and thus the least fecund at the temperature assumed to be optimal (Fig. 1B). Therefore, the specialization showed some signs of the *jack of all trades, master of none* (Huey and Hertz, 1984) pattern; clones specialized to warm performed either extremely poorly at the lowest temperature (clone Warm2) or extremely well at the highest temperature (clone Warm1), while two of the medium clones, with the exception of the largest one, performed relatively well across all of the regimes. The interpretation of cold preference is not straightforward because we had only one clone previously acclimated to low temperature. However, our results on the population growth rate (including the thermal extension from a pilot study provided in the supplement) as well as the data on fecundity together suggest that our cold-preferring clone may also be perceived as a specialist.

The results showed that 30 °C was optimal (= gave the highest mean performance) for two clones, Warm1 and Int2. However, in the case of the remaining three clones, the difference in performance between 25 °C and 30 °C was relatively small, and in each case, the value achieved at 30 °C was much higher than that at 20 °C (Fig. 2A). Therefore, the discrepancy between 30 °C and the real optimal temperature for all six clones should not considerably affect the results on fecundity, which we assumed to measure at the optimum.

Our results provide a new scientific perspective on the issue of the performance of generalists *vs*. specialists: specialists allocate more in reproduction and are therefore more fecund than generalists at optimal temperatures, while generalists display stronger plastic size response to temperature (the temperature-size rule). This result shows a possible empirical solution to the distinction between specialists and generalists first introduced by Levins (1968): specialists allocate more in reproduction at optimal temperatures, while generalists invest in plastic responses across environments.

## Acknowledgments

We are grateful to Maurizio Manca for helpful comments on the previous version of the text. This work was supported by the National Science Centre Poland (OPUS 2015/19/B/NZ8/01948), the National Center for Research and Development (GEKON1/O3/214361/8/2014) and the Jagiellonian University (DS/INoS/757/2018). The manuscript was edited by American Journal Experts. The authors declare no conflict of interest.

---

**Figure S1.**
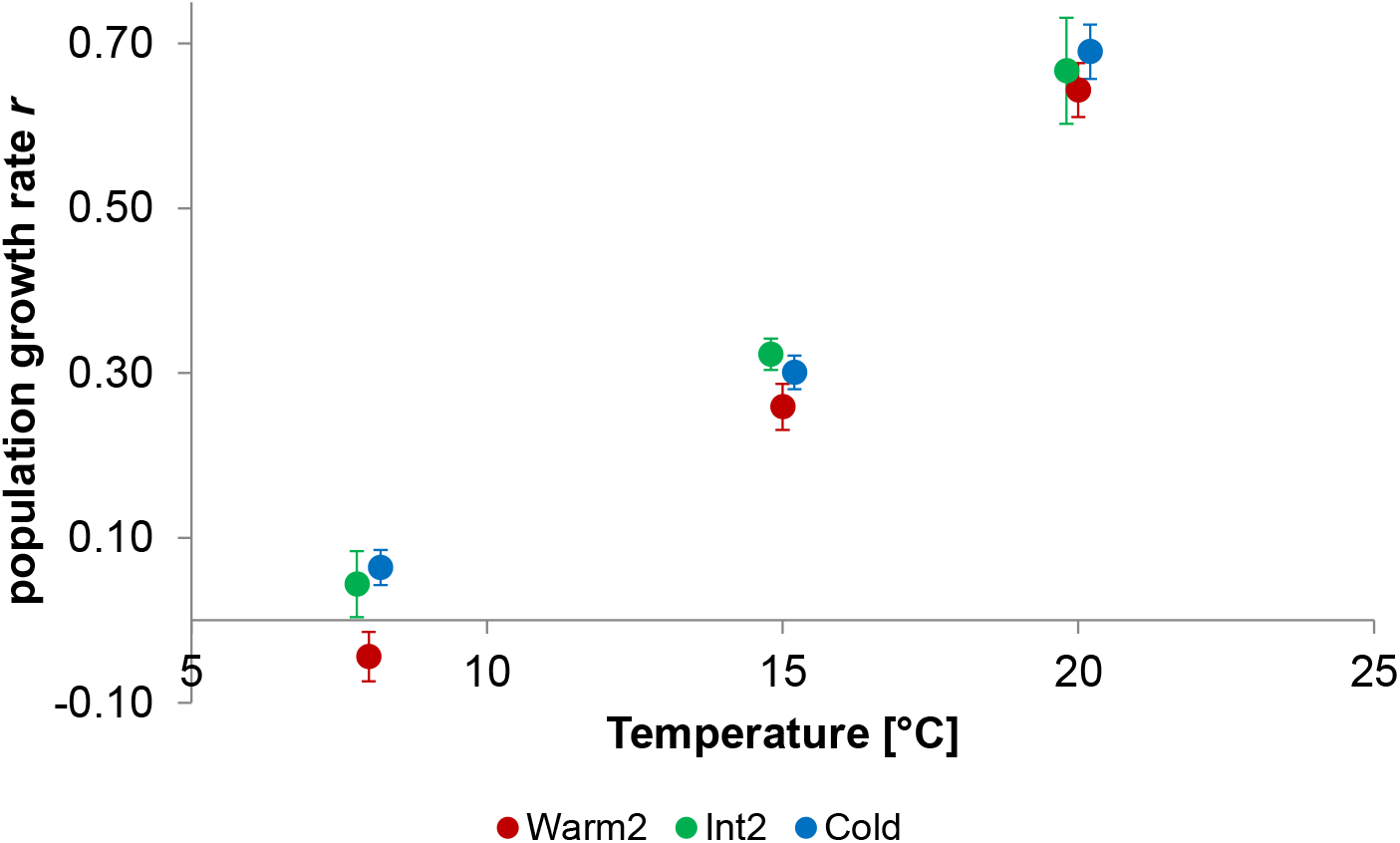
Pilot results (J. Starzycka, unpublished) for the three clones out of six investigated in this report. The population growth rate was estimated at 8, 15 and 20 °C, in the 10 three-day periods, for eight replicates per clone. The data show the mean value from nine estimates (excluding the first, 0-3 days estimate), across the replicates. Mean ± SD

